# Long mounting with repeated copulations in *Zygogramma bicolorata*: A test of adaptive significance of a curious mating behaviour

**DOI:** 10.1101/2024.01.10.574980

**Authors:** Rabi Sankar Pal, Bodhisatta Nandy

## Abstract

Long matings are abundant in insects despite the range of the costs involved. The causes and consequences of the evolution of long matings remain an interesting problem for behavioural ecologists. We studied the Parthenium beetle (*Zygogramma bicolorata*), which is known to show extraordinarily long copulations. Our detailed investigation showed, for the first time, that the long mounting in this species entails repeated copulation. Each of these copulations is preceded and succeeded by a leg-rubbing behaviour. We found the same characteristic pattern in laboratory and semi-natural conditions. We conducted a series of interrupted mounting assays to examine the fitness consequences of the duration of mounting. We tested multiple hypotheses regarding the transfer of sperm and non-sperm components, and the function of leg rubbing in modulating the female copulatory behaviour. Sperm transfer and fertility did not exhibit a linear increase with the number of copulations. There was also no evidence of nutrient transfer by the males. In contrast, our experiments reveal significant cost of long mounting in these beetles. We showed female remating behaviour can be modulated by the long previous mounting. Our data also suggest that leg rubbing may reduce female resistance to mating. While the long mating possibly serves as copulatory mate guarding, the males may increase their competitive ability through modulating female remating. We also propose leg rubbing to be a copulatory courtship behaviour. Our study provides preliminary insights into the evolution of long mating in such systems and necessitates further research on sexual selection.

## Introduction

Insects show enormous variability in their mating behaviour, including the duration for which matings last (Simmons, 2001). The latter is known to be an important trait that includes the act of transferring sperm, seminal fluid components, including nutrients, and represents the time and effort invested by a male in mating with a female (Parker, 1974; Mazzi et al., 2009; Bretman et al., 2010). Usually, a short copulation can potentially serve the purpose of reproduction *per se* as the transfer of sperm is rarely a time consuming process (Gilchrist & Partridge, 2000, Katsuki & Miyatake, 2009, Lachmann, 2013). However, in many species copulations are long, and can show significant variation, both within and across species (Birkhead & Moller, 1992; Elgar, 1995; Field et al., 1999; Córdoba-Aguilar et al., 2009; Rull et al., 2017). It may range from a few seconds to several days (Labitte, 1919; Clements & Kerkut, 1963; Cueva Del Castillo et al., 1999; Haubruge et al., 1999; Dar et al., 2021). For example, copulation in the yellow fever mosquito *Aedes aegypti* persists for only 13Lseconds (Spielman, 1964), whereas Indian stick insect *Necroscia sparaxes* pairs copulate for up to 79 days (Sivinski, 1978). In some species, like the southern green stink bug *Nezara viridula,* copulation duration may vary from five minutes to 14 days (Mitchell & Mau, 1969; Harris & Todd, 1980; McLain, 1980). What generates and maintains such extreme variation in copulation duration is an open question in behavioural ecology.

Long and extraordinarily long matings are abundant across taxa (e.g., Wickler & Seibt, 1985; Dickinson, 1986; Stockley, 1997; Harari et al., 1999; Schöfl & Taborsky, 2002), and is particularly challenging to explain because of the time and energy costs associated with it. The evolutionary cause for the evolution of long mating is often attributed to male adaptation to the risk of sperm competition (Simmons, 2001). For long mating to evolve, the benefits accrued to the males in terms of paternity success should outweigh the cost of losing additional mating and foraging opportunities, increased risk of predation and injury, assuming the mating duration is under male control (Parker, 1970; Parker, 1974; Thornhill & Alcock, 1983; Yamamura 1986; Alcock, 1994).

Sperm competition theories suggest that transferring numerous sperm may help “numerically overwhelm” and/or displace the rival males’ sperm (Dickinson, 1986; Parker & Simmons, 1991; Simmons & Parker, 1992). Males transferring a larger volume of ejaculate through long copulations tend to have higher reproductive success (Simmons, 1987; Wedell, 1991; Sakaluk & Eggert, 1996). Such advantage could partly be due to higher number of sperm transferred. In addition, males are known to transfer non-sperm components of the ejaculate, such as, seminal fluid proteins and peptides, that may aid in fertility and competitive ability (Gillott, 2003; Poiani, 2006). Seminal fluid, a cocktail of lipids, proteins, carbohydrates, salts, hormones, nucleic acids, and vitamins, comprises secretions from accessory glands, the ejaculatory duct, and the ejaculatory bulb (Chapman, 2001; Scolari et al., 2021). Seminal fluid proteins and especially accessory gland proteins (Acps) can modulate a range of behavioural and physiological traits in females both in the short and long term (Chen, 1984; Chapman et al., 1995; Cordero, 1995; Ravi Ram & Wolfner, 2007). This includes sperm storage, oogenesis, ovulation, female receptivity, and longevity (reviewed in Chapman, 2001). Acps are also used in forming mating plugs to prevent sperm loss and/or further mating attempts (Lung & Wolfner, 2001). Acps transferred by the males can potentially be beneficial for them, but deleterious to the females in terms of decreased reproductive success and longevity (Rice, 1992; Arnqvist & Rowe, 1995; Chapman et al., 1995; Rice, 1996; Civetta & Clark, 2000). Mating may also involve copulatory/post-copulatory courtship aimed at increasing paternity success (Eberhard, 1991; Eberhard, 1994; Eberhard, 1996), especially if there is a substantial scope of cryptic female choice. However, not much is known about such copulatory courtship and its fitness consequences to the males and females.

Additionally, long copulations can also be beneficial for the females. The nutrients transferred during copulation may increase females’ longevity, lifetime fecundity, or contribute to egg provisioning (Sakaluk & Cade, 1980; Gwynne, 1984; Boucher & Huignard, 1987; Bultin et al., 1987; Simmons, 1988; Wiklund et al., 1993; Karlsson, 1998; Wiklund et al., 1998; South & Lewis, 2012). However, except in some of these special cases, mating, especially long matings, appears to be detrimental to the females across taxa (Edvardsson & Tregenza, 2005; Mazzi et al., 2009; Edward et al., 2015).

Therefore, notwithstanding the advances in our understanding of sexual selection and mate choice, long matings continue to be a puzzle with ample scopes of empirical investigations. Here, we investigated Parthenium beetles, *Zygogramma bicolorata*, a species known to show extraordinarily long matings (Afaq & Omkar, 2017, Bhaisare et al., 2021; Bhaisare & Chaudhary, 2023). Afaq & Omkar (2017) found long mating to be associated with increased adult reproductive performance and improved offspring life-history traits. In another study, the prolonged mating in this beetle did not contribute to the increase in fecundity and hatchability beyond certain duration, leading the authors to conclude that such a behaviour to be a form of mate guarding (Bhaisare et al., 2021).

We revisited this problem and described the mating behaviour, specifically probing the duration for which a pair appears to be in copula. We show that the long copulation duration mentioned in the previous studies is actually the period of mating association, which includes several short, repeated rounds of copulation interspersed with an intriguing set of inter-copulatory behaviours. We first characterized the mating behaviour in the laboratory set-up. Additionally, we have carried out observations in semi-natural conditions to rule out the possibility of these behaviours being artefacts of static laboratory conditions/experimental environment. We further examined the fitness consequences of variation in the number of copulations per mating association to eventually explain the ultimate level explanation for this behaviour. We investigated the transfer of sperm and non-sperm components that could play a major role in shaping the reproductive ecology in systems showing long mating associations. Finally, we looked into male inter-copulatory behaviours explaining its possible role in influencing female behaviour during repeated copulations.

## Methods

*Zygogramma bicolorata* Pallister (Coleoptera: Chrysomelidae) is a phytophagous beetle of Neotropic origin (Ganga Visalakshy et al., 2008). Presently widespread and abundant in most areas of India, it was originally introduced in India in the 1980s for the biological control of invasive *Parthenium hysterophorus* (Jayanth, 1987). These beetles can complete their life cycle only on a few species of *Parthenium* and *Ambrosia* (McFadyen & McClay, 1981). They are promiscuous and present in the wild primarily in a female-biased sex ratio (Hasan & Ansari, 2016a). There is little sexual dimorphism except that the females are larger (Afaq & Omkar, 2013). Eggs are laid on the leaves singly or in groups (Dhileepan et al., 2019). Larvae feed voraciously on the leaves and inflorescence, undergo four instars, and pupate under the soil (Dhileepan et al., 2018). The life cycle in nature is completed in 4–8 weeks, depending on environmental parameters (Dhileepan & Wilmot Senaratne, 2009; Sushilkumar & Ray, 2011; Hasan & Ansari, 2016b). These beetles may undergo diapause for up to six months in a year (Dhileepan & Wilmot Senaratne, 2009).

### Founding population and laboratory maintenance

A laboratory population was established with 30 females and 30 males collected in January 2021 from six locations of Berhampur, Odisha, India (19.3150°N, 84.7941°E). The wild-collected beetles were given 48 hours to acclimate to the laboratory environment. Five females and five males were kept together in a transparent plastic container (henceforth adult box) of volume 500 ml (15 cm diameter, 8 cm height) at 26 (±2) °C under continuous exposure to light. Each adult box was supplied with freshly collected *ad libitum Parthenium* leaves. The petioles of the leaves were placed in small plastic tubes filled with tap water to keep the leaves fresh. Following 48 hours, fresh leaves were introduced, and eggs laid on this petiole in the next 2–3 days were collected to start the first laboratory generation. Eggs started to hatch day 4 onwards. Newly hatched larvae were kept in aerated petri dishes supplied with freshly collected leaves, which are replaced on alternate days. Petioles of the leaves were always wrapped in wet tissue paper to keep them hydrated. As the larvae grew, juvenile density in these plates was controlled by redistributing larvae in additional plates to avoid the adverse effect of competition between larvae. This was done by keeping the number of larvae per petri dish fixed for a given instar stage. Twenty, ten, eight, and five individuals of first, second, third, and fourth instar larvae, respectively, were held in a single petri dish with leaf. On the day 17 from oviposition, fourth instar larvae were transferred to the pupation boxes (plastic containers of 500 ml volume, filled with moist sand (up to three-fourth the volume) and leaves on top (for larvae to forage). Once they emerged as adults from the sand, males and females were identified under the microscope by their last abdominal sternite (McClay, 1980). On eclosion, virgin adult beetles were collected at 24-hour intervals, and were held in adult boxes. 8–10 individuals of single sex were kept in each adult box. Eclosion into adults from eggs took 27–33 days. The beetles reached sexual maturity about day 11 post-eclosion. On day 11, the sexes were combined in groups of five of each sex in the adult boxes. Eggs laid in the next 2–3 days constituted the next generation.

Same maintenance protocol was followed each generation. Population size of the laboratory stock was roughly 350–400 individuals per generation. To reduce the effect of inbreeding, the laboratory population was supplemented with 25 males and 25 females collected from the natural population of Berhampur in the fourth laboratory generation.

### Mating behaviour in laboratory conditions

On the 11^th^ day post-eclosion, beetles were kept individually in glass vials (25 mm diameter, 96 mm height) with leaves for 24 hours for pre-conditioning prior to the observation. A virgin female and an unmated male were picked randomly and paired in each vial the next day. Small pieces of *Parthenium* leaves were supplied inside the vial prior to the pairing. Each pair was given four hours to initiate mating. A mating association initiated when a male dorsally climbed the female, and continued till the pair dismounted. The pairs that did not start mating during these four hours were removed and not observed further.

During a mounting episode a pair usually goes through a series of short copulations. Each copulation was defined as the insertion of the male aedeagus into the female’s genital opening. Release time, mounting time, beginning and end time of each copulation were recorded closest to a minute. Mating association will hereon be referred to as mounting.

We also observed two relatively high-frequency behaviour - leg rubbing and kicking. We conducted a separate assay to quantify these two components of behaviour aiming to assess their role as copulatory courtship.

A total of 48 successful matings were observed by multiple observers over five days to accommodate the logistic limitations. The observations started around 10 a.m. every day and continued until the pairs finished mounting. One observer observed a maximum of three pairs at any given time point. All the mating vials were kept under the light during observation.

Mounting latency (the time male takes to dorsally mount the female after release into the mating arena), copulation latency (the time between mounting and the beginning of the first copulation), copulation duration (the duration of a single intromission), inter-copulatory latency (The period between two subsequent copulations), and mounting duration (the time between mounting and dismounting of a pair that includes repeated copulation) were quantified from the observations.

We observed that an entire mounting duration of a pair of *Z. bicolorata* was a series of copulations (see Results section for quantitative description). To assess the adaptive significance of long mounting seen in this species, we carried out a series of interrupted mounting assays and measured the fitness consequences of the experimental perturbation of mounting (Schematically outlined in Figure 1).

**Figure 1:**
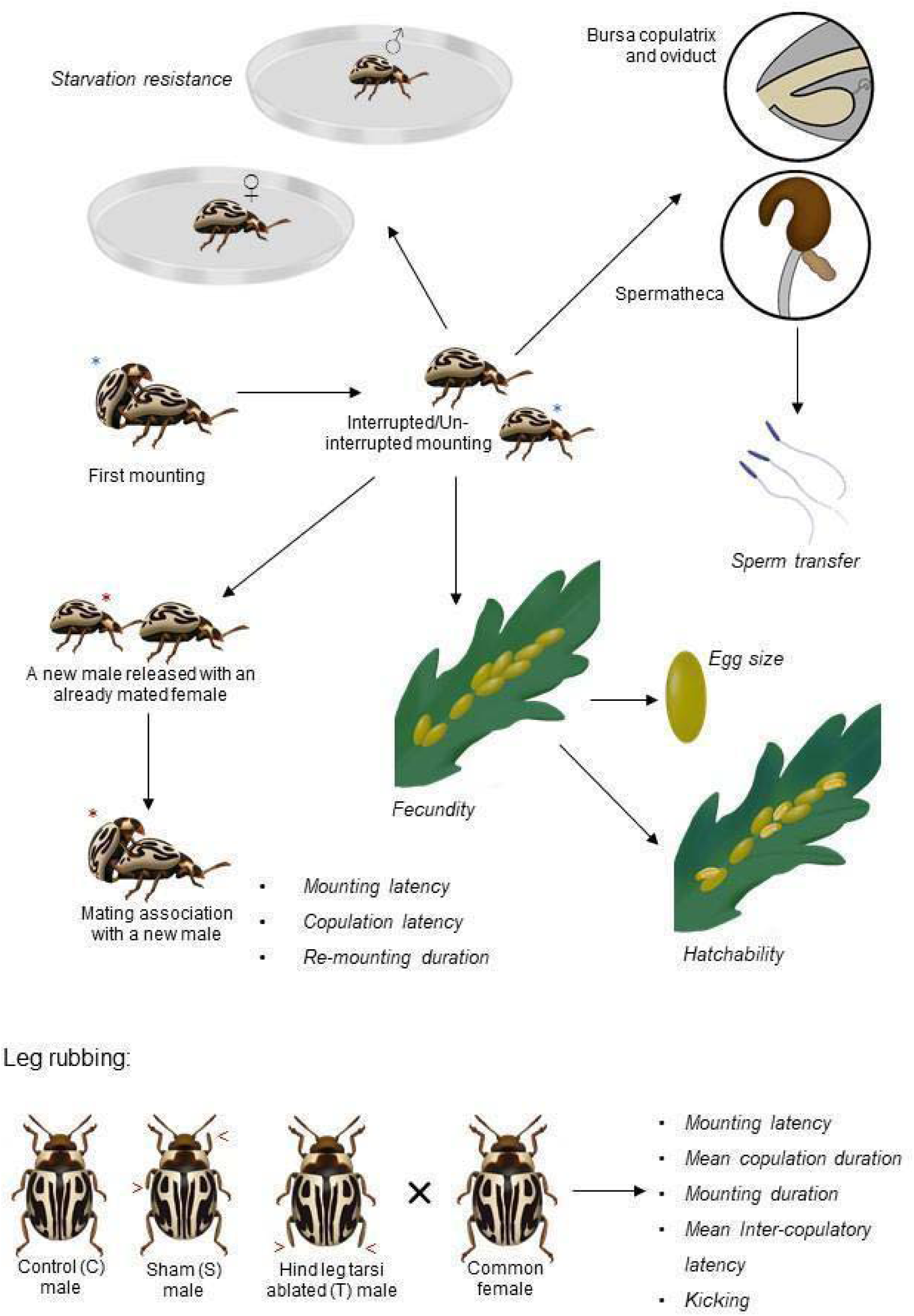
Schematic representation of the experiments done in this study investigating the mating behaviour of *Zygogramma bicolorata*.

### Experiment 1: Sperm transfer and fertility

The first set of interrupted mounting assays were done in the fourth laboratory generation. This experiment was aimed at testing the theory that long mounting and repeated copulations are necessary to ensure sperm transfer, and fertility. We investigated the effect of experimental truncation of mounting duration on sperm transfer, female fecundity, and hatchability.

A mating vial was set up by combining a 12-day old virgin female and a male in a glass vial. A total of 180 vials were set up. Out of the pairs that initiated copulation, 110 vials were randomly assigned to four mounting interruption treatment groups – C1, C15, C30, C50, and a control set. Mounting was interrupted by gently removing the male with a paintbrush following the completion of 1 (C1), 15 (C15), 30 (C30), and 50 (C50) copulations. In the control set, the pairs were allowed to complete their mounting. The interruption treatments were chosen based on Afaq & Omkar (2017) where fecundity and egg viability showed significant increase up to mating for two hours, beyond which there was little or no difference. In our experiments, C15 and C30 roughly correspond to one and two hours of mounting, respectively.

### Sperm transfer

Ten females from each group were anaethetized in mild exposure to CO_2_ and dissected within two hours of the interruption or completion of mounting, as specified by the treatment group. We used a method modified from Bloch Qazi et al. (1996) and Baer et al. (2016) for the estimation of sperm number. Spermatheca was dissected out in a lid of 0.2 ml PCR tube containing 50 µl 1X phosphate buffered saline (PBS) of pH 7.2. The spermathecal walls were ruptured with a fine forcep and stirred in PBS such that the entire sperm content was dispersed into the PBS. Similarly, the bursa copulatrix and oviducts were dissected in 25 µl PBS, severed in multiple places, and agitated gently to release the sperm content into PBS. The samples were vortexed gently for 10 seconds. An equal volume of DAPI working solution (1 µg/ml) was mixed with the sperm solution and incubated in the dark for 20 minutes. DAPI-stained sperm heads were counted in the improved Neubauer chamber under 400x magnification with a DAPI filter in Zeiss Scope.A1 fluorescence microscope. Each sample from the spermatheca and bursa copulatrix-oviduct was counted twice in four 1 mm^2^ grids of the Neubauer chamber. Since the volume of each 1 mm^2^ grid is 0.1 µl, total sperm count in the spermatheca and bursa copulatrix-oviducts was obtained by multiplying the mean number of sperm in each 1 mm^2^ grid with 1000 and 500, respectively.

### Fecundity and hatchability

Our pilot experiment (data not shown) indicated that the cumulative lifetime fecundity of the singly mated females experienced a sharp increase until about day 60 post-mating before it levelled off. To understand the fitness consequences associated with long mating, the fecundity and hatchability of the females from the above-mentioned treatments were measured in a five-day interval starting from day 1 to day 61 post-mating. Each treatment had 12 mated females. Any effect of the number of copulations on cumulative fecundity and mean hatchability was examined.

### Experiment 2: Potential role in modulation of female remating

Insect males often pass on accessory gland proteins (product of accessory glands of the male reproductive system) that aid in paternity defence through the modulation of female remating tendency and duration of remating (Leopold, 1976; Wolfner, 1997; Radhakrishnan & Taylor, 2007; Yang et al., 2009; Wigby et al., 2020). Long mounting in this species could potentially be a putative paternity assurance strategy. We tested this theory by measuring different components of remating behaviour of females subjected to interrupted mounting treatments. 12-day old virgin males and females from the sixth generation following laboratory domestication were paired in petri dishes with leaves following a 24-hour acclimation period. The copulating pairs were randomly assigned to three treatments – (a) 30 copulations (C30); (b) 50 copulations (C50), and (c) Control, where mounting pairs were allowed to naturally dismount. Sperm storage in females can, in principle, modulate the receptivity of the females (Gromko et al., 1984; Letsinger & Gromko, 1985; Oberhauser, 1989; Wedell, 1993). Therefore, C30 and C50 were selected for the assay. Compared to controls, these two interrupted mounting treatments were found to have no significant difference in the sperm transfer (see Results). Following interruption/completion of the mounting, the mated males were removed, and a new unmated male was released immediately with the already mated females. Remating assay started with 21 mated females in C30 and C50, and 20 in the control group. Mounting latency, copulation latency, and the duration of the second mounting (i.e. re-mounting) were noted. Given the design of the assay, a systematic difference in the time (of the day) when the target behaviour (i.e., remating) was observed, could not be avoided. Depending on the treatment, remating observation was done at different times of the day – on an average, first, C30, then C50, and finally the controls. We conducted a separate assay to rule out the possibility of such systematic difference across treatment was not a confounding factor. The details of this assay can be found in the supplementary information. Briefly, we showed that different components of mating behaviour, including mounting duration, was not affected by the time of day (see Supplementary information). Hence, we argue that time of the day was unlikely to be a confounding factor in our Experiment 2.

### Experiment 3: Potential nutrient transfer

Mating in many insects, including many beetles, entail nutrient transfer from the male to the female (Vahed, 1998; Rooney & Lewis, 1999; Savalli & Fox, 1999; Takakura, 2004). Nutrient received during mating may increase female survival, fecundity and/or egg size that may improve progeny fitness (Parker & Simmons,1989; Boggs, 1990; Fox & Czesak, 2000; Giron & Casas, 2003). Hence, it is possible that long mating in *Z. bicolorata* represents such nutrient transfer. If so, interrupted mounting should result in reduced fecundity and/or egg size and/or decreased female starvation resistance. Fecundity data was collected as a part of Experiment 1. We tested the latter two predictions by carrying out two separate interrupted mounting experiments similar to Experiment 1 described above.

The egg size assay was conducted by assigning five sets of randomly chosen ten females to five treatment groups (C1, C15, C30, C50, and control). The experimental females were kept individually in petri dishes on completion or interruption of the mounting. Fresh Parthenium leaves were supplied on alternate days. The females deposited their eggs on the leaves. Approximately 30 eggs from each treatment were harvested for size measurement. Egg size was measured on the day 1, 5, 10, followed by a 10 day interval until day 60 post-mating. The first three samplings were done in shorter intervals to inspect any short-term effect of the number of copula on egg size. Approximately equal numbers of eggs were sampled from each of the lobes and midrib regions of the parthenium leaf. Eggs were placed on a Petri dish and images were captured in Zeiss Stemi 508 microscope fitted with Zeiss Axiocam ERc5s using Zen 2.3 (blue edition) software. Egg size was measured in ImageJ (Version 1.53k) (Schneider et al., 2012) to the nearest 0.1 µm.

A separate set of beetles were subjected to experimental treatments followed by starvation (and desiccation) survival assay. Two interrupted mounting treatments-C1 and C30 (see above) were used in this assay. Virgins and pairs that naturally dismounted were taken as negative and positive controls, respectively. Once a mounting was interrupted/completed, both males and females were transferred individually to Petri dishes without leaf. The number of days each individual survived were counted. Starvation survival time (time to death under starved condition) was considered a measure of starvation resistance. While difference in female survival under starvation in this assay could be indicative of nutrient transfer during mating, male starvation resistance can indicate the cost of mating. C1, C30, and complete mating treatments had 25, 26, and 24 males and females, respectively, along with 25 unmated males and 27 virgin females. The assays were conducted in the eighth laboratory generation.

### Experiment 4: Leg rubbing as copulatory courtship

In the ninth generation, five-day-old males were divided randomly into three groups. Tarsi of the two hind legs were ablated in treatment (T) group males. In the control (C) group of males, no ablation was performed. Additionally, a sham treatment (S) was used, in which tarsal ablation was carried out in one of the fore and mid legs, alternatively on the left or right side. The ablations were performed on day 5 post-eclosion under CO_2_ anaesthesia. The males and females were paired on post-eclosion day 12 under light. The pairs were observed for a period of 20 minutes in roughly one-hour intervals from the beginning of the copulations until the end of the mounting, or up to eight observation periods, whichever is earlier. One observer observed only one pair at any point in time. The time of release, mount, and beginning and end of each copulation were noted closest to a second, in addition to the number of times females kicked the males during observation periods. The T, C, and S groups had observations from 12, 11, and 11 pairs, respectively.

Edvardsson & Arnqvist (2000) warned that phenotypic manipulation could generate behavioural artefacts and that manipulation should not interfere with other normal behaviours. We did not observe any definite alteration in their behaviour in their normal ecology, including foraging and movement. The males tried to perform leg rubbing before and after each copulation in the usual manner, even after tarsal ablation.

### Ethical note

This work followed the ASAB/ABS Guidelines for the ethical treatment of nonhuman animals in research. Dissections and tarsal ablations were performed under CO_2_ anaesthesia. The males were monitored following ablation to ensure normal behaviour. Animals’ welfare was diligently prioritized during housing, handling and experiments. Beetles used in this study do not come under the purview of ethical approval.

### Statistical Analyses

All the analyses were performed in R version 4.2.0 (R Core Team, 2022). In all the analyses, data points outside the 1.5 × interquartile range were removed as outliers. The normality of the residuals of the model, wherever required, were tested by the Shapiro-Wilk test. The *lm* function was used to fit Linear models to data when the residuals of the model followed normal distribution. Otherwise, Generalized linear models (GLM) and Generalized linear mixed models (GLMM) were implemented using the `lme4` (Bates et al., 2015) package. Canonical link functions were employed for particular error structures specified in the models. Interruption treatments and ablation treatments were modelled as fixed factors. Overdispersion and zero inflation were checked where applicable, using the `performance` package (Lüdecke et al., 2021). To obtain *p* values, `Anova` function from the `car` package was used (Fox & Weisberg, 2019). Further, `emmeans` package was used for pairwise contrasts (Lenth, 2022). Plots were created using the `ggplot2` package (Wickham, 2016).

Comparison between first, last and mean copulation durations were done using Wilcoxon signed-rank test. The number of sperm transferred (Experiment 1) and egg size (Experiment 3) were analyzed using GLM with Gamma error distribution. The cumulative fecundity (Experiment 1) and the re-mounting duration (Experiment 2) were fit into linear models. Effect of interruption treatment on mean hatchability was evaluated using GLM with the binomial error structure following correction for overdispersion by incorporating an observation level random effect into the model (Harrison, 2014). GLM with negative binomial distribution were fitted to the mounting latency in Experiment 2 and mounting latency data in Experiment 4 using the `MASS` package to correct for overdispersion (Venables & Ripley, 2002). To account for zero inflation, we used zero-inflated negative binomial regression to analyze the copulation latency data in Experiment 2 using the `pscl` package (Jackman, 2020).

Starvation resistance was analyzed using Cox Proportional Hazard models with interruption treatment and sex as fixed factors using the `survival` package (Therneu, 2022). To plot survival curves `ggquickeda` (Mouksassi et al., 2022), and `survminer` (Kassambara et al., 2021) packages were used. Since sex and treatment×sex, both had insignificant effect on survival, we removed sex from the model while checking for the multiple comparisons. Multiple comparisons of survival curves were obtained using the `survminer` package following *p*-value adjustment using Benjamini-Hochberg procedure (Benjamini & Hochberg, 1995).

The effects of ablation treatment on mounting duration, kicks per minute, and mean copulation latency (Experiment 4) were tested by fitting linear models to the data. Mean copulation duration in Experiment 4 was analyzed using the log-normal regression to meet the assumption of normality.

## Results

In the laboratory setup, males often approached the female from behind and mounted dorsally on female elytra. The males took ∼39 minutes (mean ±SE: 38.64 ±5.50) from release to mount in the laboratory setup. In the semi-natural setup, once a male-female pair was released into the setup, the male either quickly mounted the female, or moved apart. We observed the former, and found that in such cases, mounting latency was approximately 1 minute (mean ±SE: 1.00 ± 0.22). No pre-copulatory courtship behaviour was distinctly recognizable. Females exhibited a sidewise shaking behaviour that often deterred the approaching male and sometimes dislodged the one that already mounted. When mounted, tarsi of the pro- and meso-thoracic legs rest on the female elytra while tarsi of the hind legs rest on the edge of the elytra. Aedeagus of the male may be extruded during mounting. The copulation generally started in less than a minute since mounting in both laboratory (mean ±SE: 0.98 ± 0.26 minute) and semi-natural (mean ±SE: 0.35 ± 0.12 minute) condition.

A single long mounting appeared to consist of a series of copulations. Most of the copulations took 3 minutes in laboratory observation and 2 minutes in semi-natural condition observation. The two consecutive copulations were separated by a latent period of 1 minute both in lab and semi-natural observation. Each copulation was preceded and followed by leg rubbing behaviour by the male in which the male rubbed the lateral edges of the last abdominal segment with the tarsi of their hind legs. The behaviours in the sequence (copulation, leg rubbing and inter-copulatory latent period) were repeated in a rhythmic manner (Figure S1). In laboratory and semi-natural setup, single mounting consisted of ∼55 (mean ±SE: 55.5 ± 5.04) and ∼57 (mean ±SE: 56.75 ± 9.24) copulations, respectively (Figure 2, Table S1). Mean copulation duration was close to 4 minutes (mean ±SE: 3.95 ± 0.16) in the laboratory condition and about 3 minutes (mean ±SE: 2.67 ± 0.11) in the semi-natural condition, whereas mean inter-copulation latency varied between 1–2 minutes (mean ±SE: 1.52 ± 0.11 minute in laboratory condition, and 1.31 ± 0.15 in semi-natural condition) (Figure 2, Table S1). Towards the end of each copulation, the aedeagus appeared to be inserted deeper into the female genital opening before retracting it. During copulation male mouthparts periodically touched the female elytra and the antennae of the male were positioned sideways. Sporadically, some females showed abdomen curling (hiding their abdomen upward inside the elytra making genital opening inaccessible to the males). Males were seen biting the mesofemur or mesotibia of the females during the inter-copulatory latent period, especially when the females did not accept copulations. Females frequently kicked the male with their hind legs during or in between copulations. The mounting continued for an extended period, ∼322 minutes (mean ±SE: 322.30 ± 24.31) in laboratory condition and ∼235 minutes (mean ±SE: 235.35 ± 35.46) in semi-natural condition (Figure 2, Table S1). In laboratory condition, mounting duration ranged from 8 to 700 minutes, whereas in semi-natural condition, it ranged from 3 to 501 minutes. In both conditions, the first and last copulations typically took longer time, compared to the mean copulation duration (Figure 2, Table S1, S2). During mating females foraged periodically while the males starved for the entire mating period.

**Figure 2:**
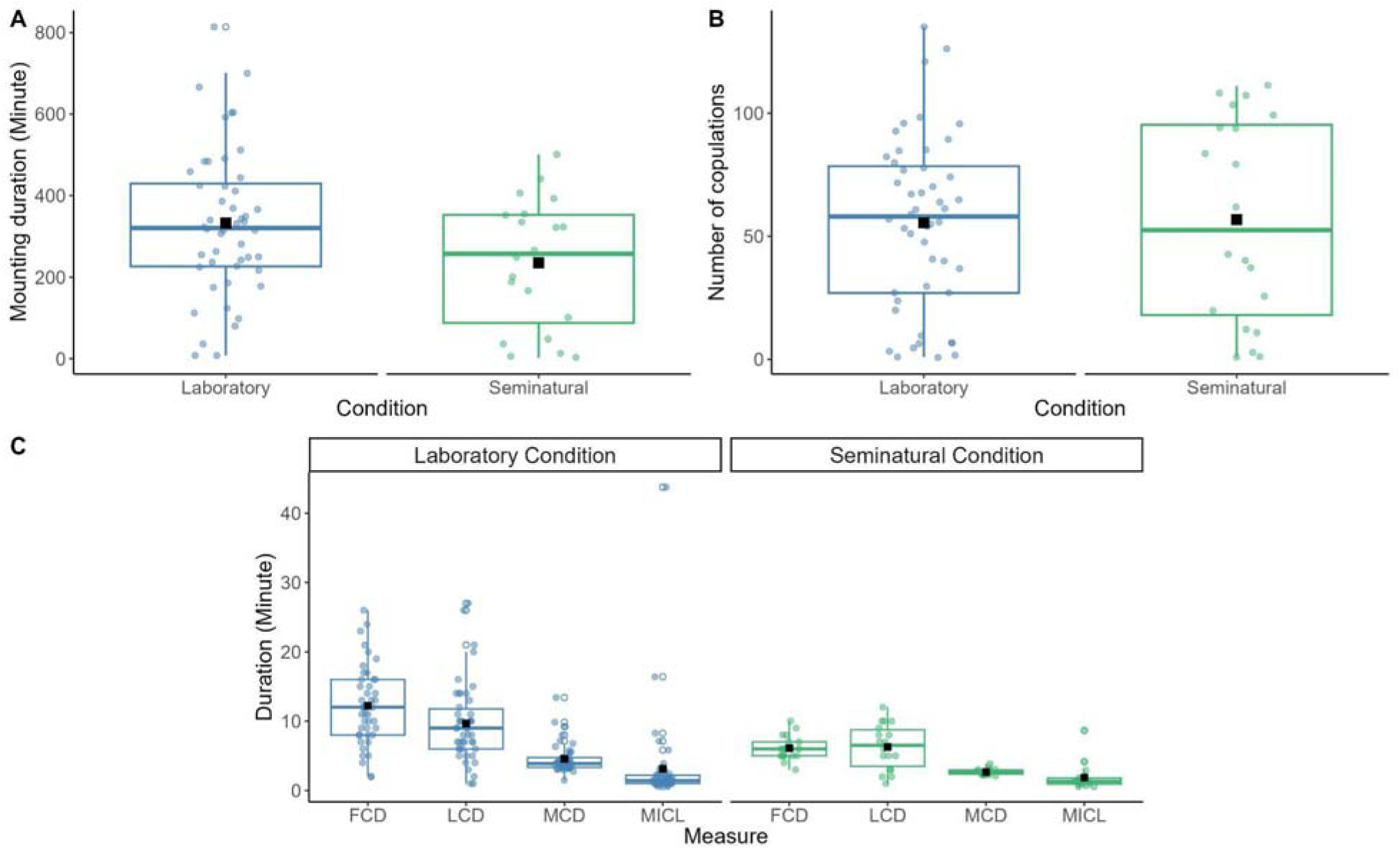
**A**) Mounting duration, **B**) number of copulations, **C**) first copulation duration (FCD), last copulation duration (LCD), mean copulation duration (MCD) and mean inter-copulation latency (MICL) of the pairs observed during the characterization of mating behviour in laboratory (n = 48) and semi-natural (n = 20) conditions. Closed and open circles indicate the individual values and outliers, respectively. Black squares represent the mean values of the observations.

### Experiment 1: Sperm transfer and fertility

Results of our analysis indicated that there was a significant effect of interruption treatment on sperm transfer, cumulative fecundity and hatchability (Figure 3, Table 1). Sperm transfer, cumulative fecundity and mean hatchability in C1 were significantly lower compared to the other treatments (Table S3–5). Whereas, the treatments other than C1 did not differ statistically from each other.

**Figure 3:**
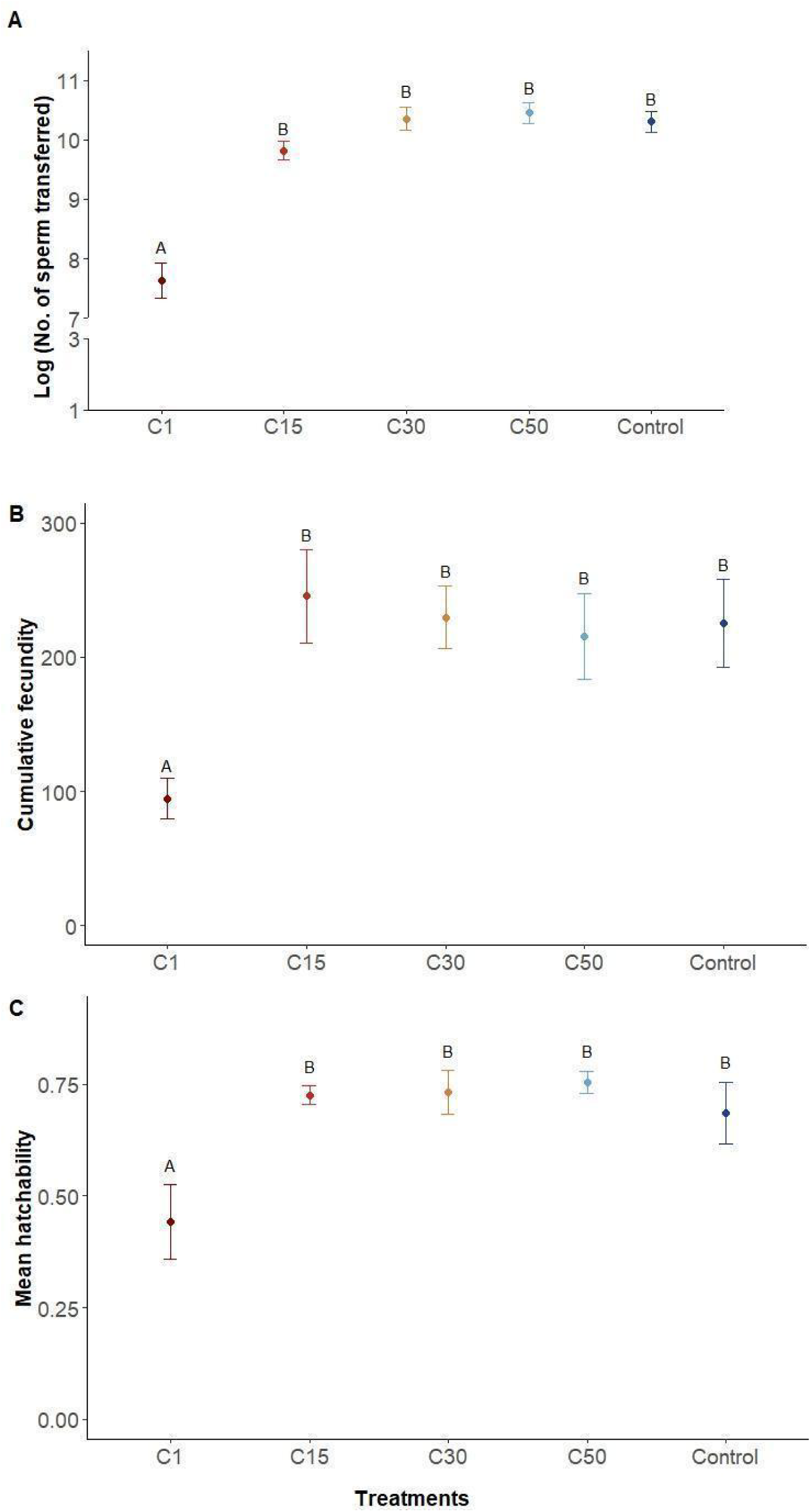
Results from interrupted mounting assay on sperm transfer, fecundity, and hatchability. Total number of sperm transferred by the males following copulation 1 (C1), 15 (C15), 30 (C30), 50 (C50) and uninterrupted mounting (Control) has been shown in logarithmic scale. The circle and the error bar represent the mean and standard error, respectively. Significant differences between treatments are marked with different letters.

**Table 1:**
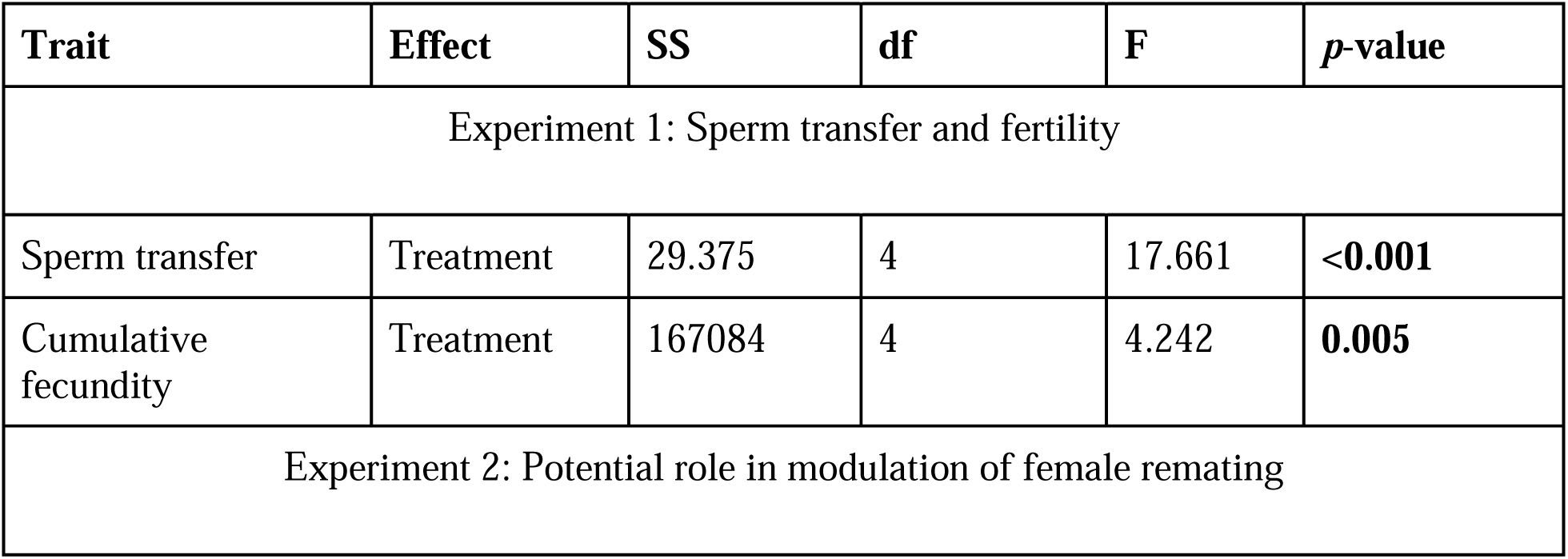
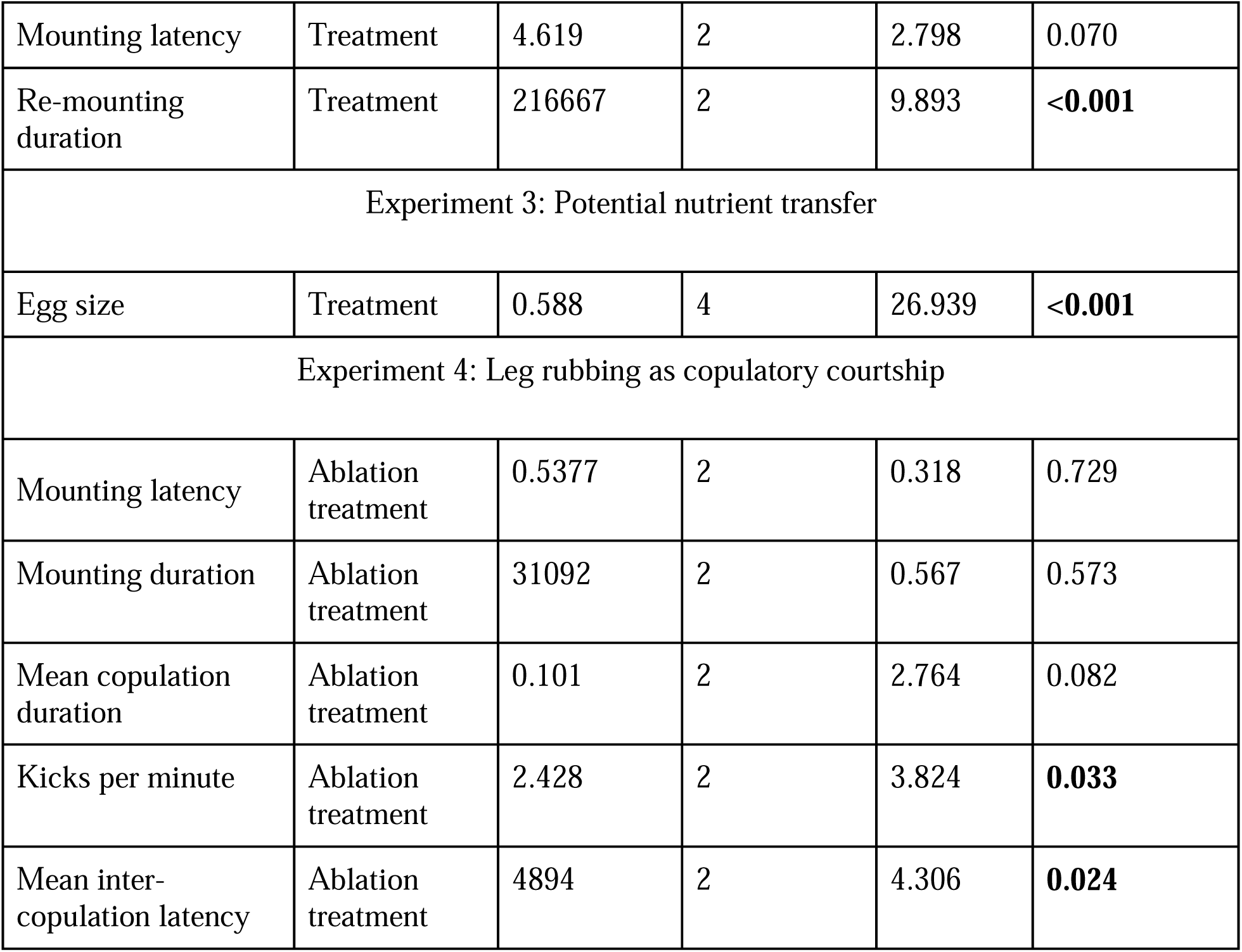
Summary of the results of various analyses. All the tests were performed considering α= 0.05. Statistically significant *p*-values are highlighted in boldface.

| Trait | Effect | SS | df | F | *p*-value |
| --- | --- | --- | --- | --- | --- |
| Experiment 1: Sperm transfer and fertility |  |  |  |  |  |
| Sperm transfer | Treatment | 29.375 | 4 | 17.661 | **<0.001** |
| Cumulative fecundity | Treatment | 167084 | 4 | 4.242 | **0.005** |
| Experiment 2: Potential role in modulation of female remating |  |  |  |  |  |
| Mounting latency | Treatment | 4.619 | 2 | 2.798 | 0.070 |
| Re-mounting duration | Treatment | 216667 | 2 | 9.893 | **<0.001** |
| Experiment 3: Potential nutrient transfer |  |  |  |  |  |
| Egg size | Treatment | 0.588 | 4 | 26.939 | **<0.001** |
| Experiment 4: Leg rubbing as copulatory courtship |  |  |  |  |  |
| Mounting latency | Ablation treatment | 0.5377 | 2 | 0.318 | 0.729 |
| Mounting duration | Ablation treatment | 31092 | 2 | 0.567 | 0.573 |
| Mean copulation duration | Ablation treatment | 0.101 | 2 | 2.764 | 0.082 |
| Kicks per minute | Ablation treatment | 2.428 | 2 | 3.824 | **0.033** |
| Mean inter-copulation latency | Ablation treatment | 4894 | 2 | 4.306 | **0.024** |

### Experiment 2: Potential role in modulation of female remating

Results of the analysis showed that there was no significant effect of treatment on mounting latency, and copulation latency (Table 1). However, the effect of treatment was found to be significant on the duration of re-mounting. Re-mounting duration of C30 and C50 females was significantly higher compared to control females (Figure 4, Table S6).

**Figure 4:**
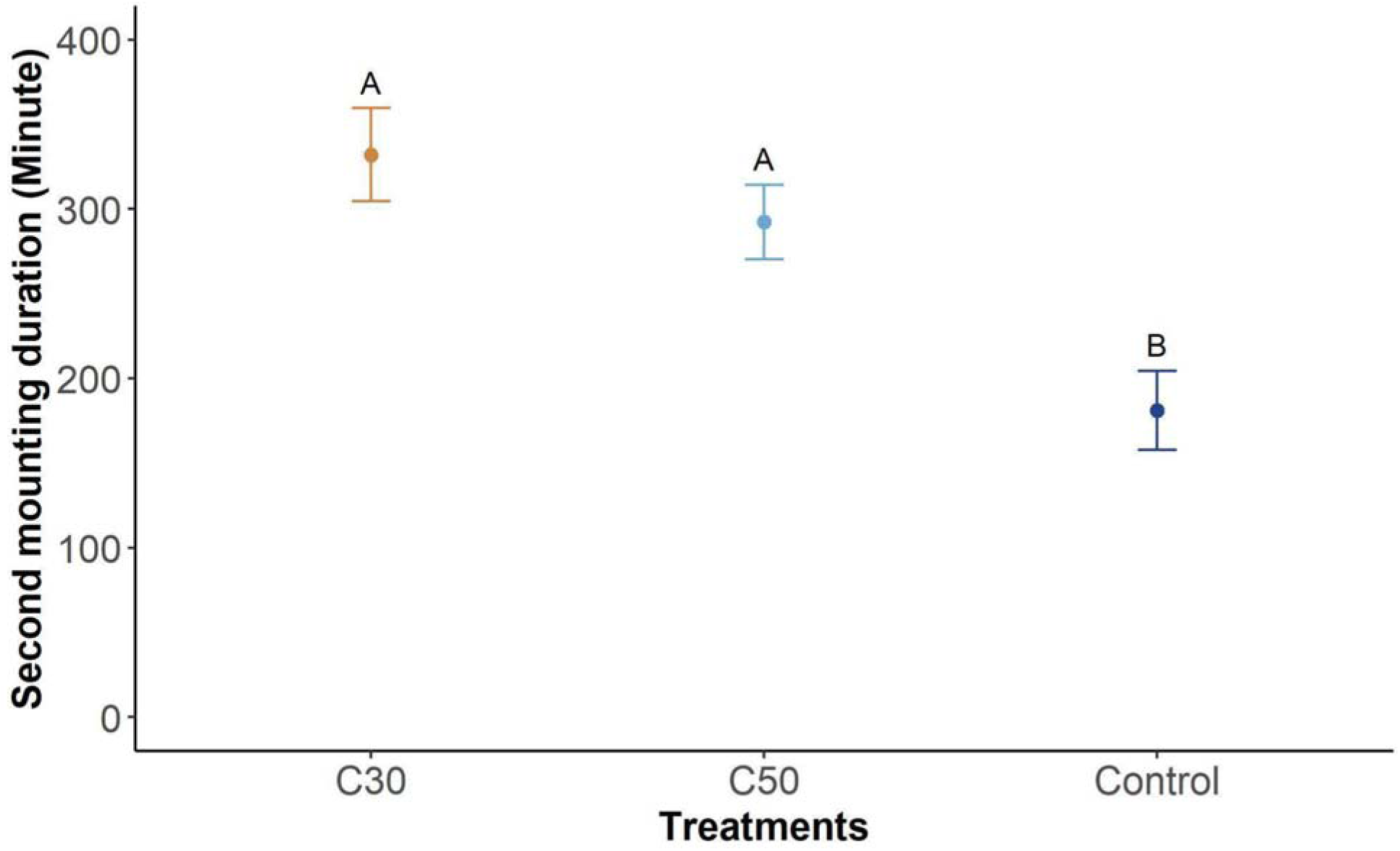
Re-mounting duration of the females that mounted till 30 (C30), 50 (C50) copulations or dismounted naturally (Control) during their first mounting. The circle and the error bar represent the mean and standard error, respectively. Treatments not sharing the same letter are statistically different.

### Experiment 3: Potential nutrient transfer

Analysis of the starvation survival time results showed a significant effect of treatment (Figure 5, Table 2). C30 (Hazard Ratio= 1.985, CI= 1.130–3.488, *p*=0.017) and Control groups (Hazard Ratio= 3.224, CI= 1.713–5.961, *p* <0.001) suffered significantly enhanced risk of mortality compared to the ‘Virgins’ (Table S7). However, no difference in the survival rates was apparent between sexes. The effect of interaction between treatment and sex was also insignificant (Table 2). Pairwise comparisons between treatments has been shown in Table S8.

**Figure 5:**
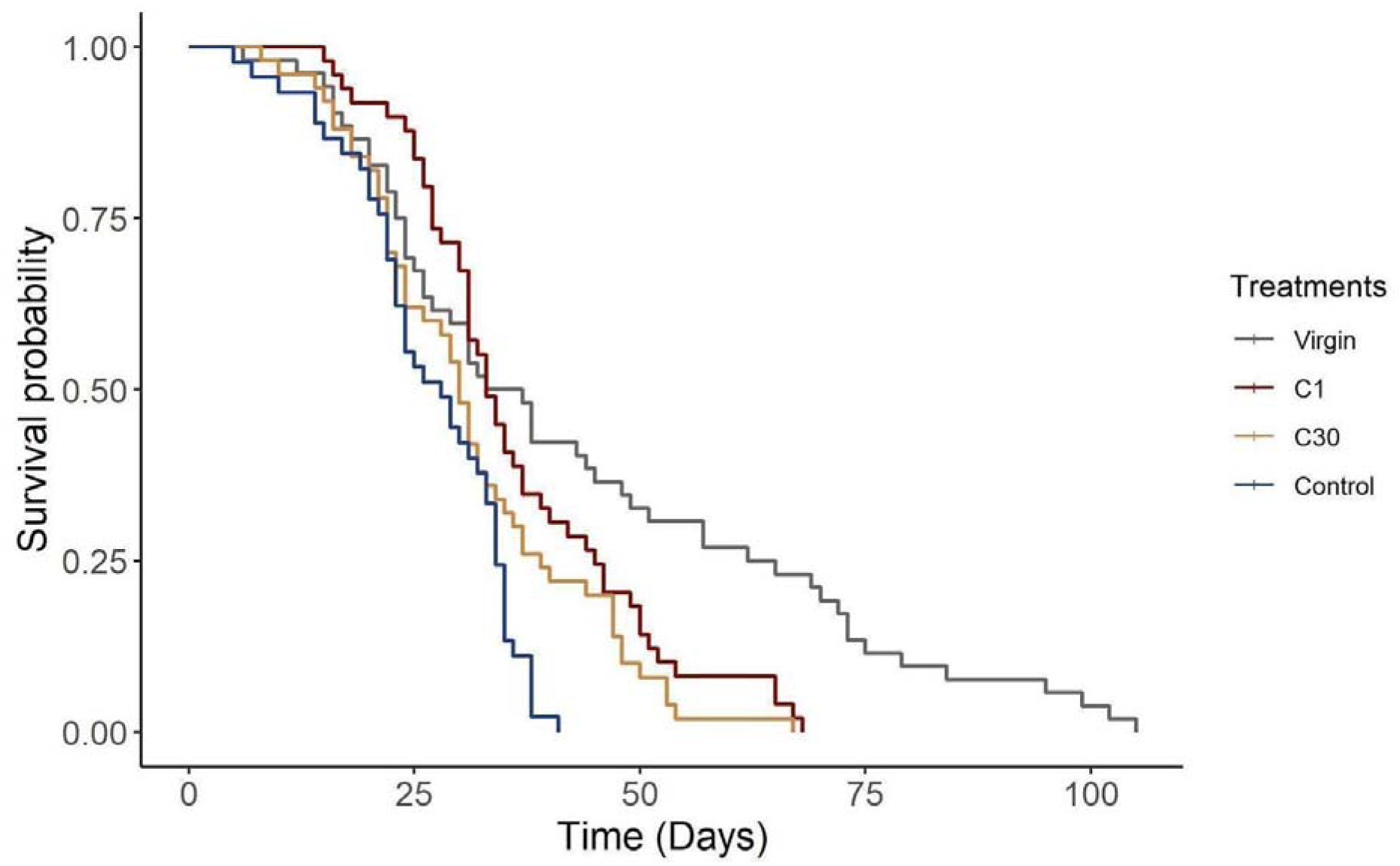
Survival plots of females and males (together) that were put into starvation as virgins or after copulation 1 (C1), 30 (C30) or naturally dismounting (Control). Survival curves were plotted using Kaplan-Meier Method. A significant effect of treatment was found.

**Table 2:** Summary of the results of various analyses. All the tests were performed considering α= 0.05. Statistically significant *p*-values are highlighted in boldface.

| Trait | Effect | Chi square | Df | $p$-value |
| --- | --- | --- | --- | --- |
| Experiment 1: Sperm transfer and fertility |  |  |  |  |
| Mean hatchability | Treatment | 25.933 | 4 | **<0.001** |
| Experiment 2: Potential role in modulation of female remating |  |  |  |  |
| Copulation latency | Treatment | 1.241 | 2 | 0.538 |
| Experiment 3: Potential nutrient transfer |  |  |  |  |
| Survival under starvation | Treatment | 14.811 | 3 | **0.002** |
|  | Sex | 0.238 | 1 | 0.625 |
| | Treatment × Sex | 1.245 | 3 | 0.742 |

Analysis of the egg size results did not indicate a pattern predicted by the nutrient transfer theory. For brevity, we have included the results and the discussion of the observations on egg size in the supplementary information.

### Experiment 4: Leg rubbing as copulatory courtship

Results of the analysis suggested that there was no significant effect of ablation treatment on mounting latency, mounting duration, and mean copulation duration (Table 1). Interestingly, ablation treatment was found to have a significant effect on the number of kicks received by the males, and the mean inter-copulation latency (Table 1). T males received significantly more kicks per minute compared to control, while S did not differ statistically from C and T (Figure 6, Table S10). The mean inter-copulation latency also displayed a similar trend (Figure 6, Table S11). T males hadd 39% higher inter-copulation latency compared to control males.

**Figure 6:**
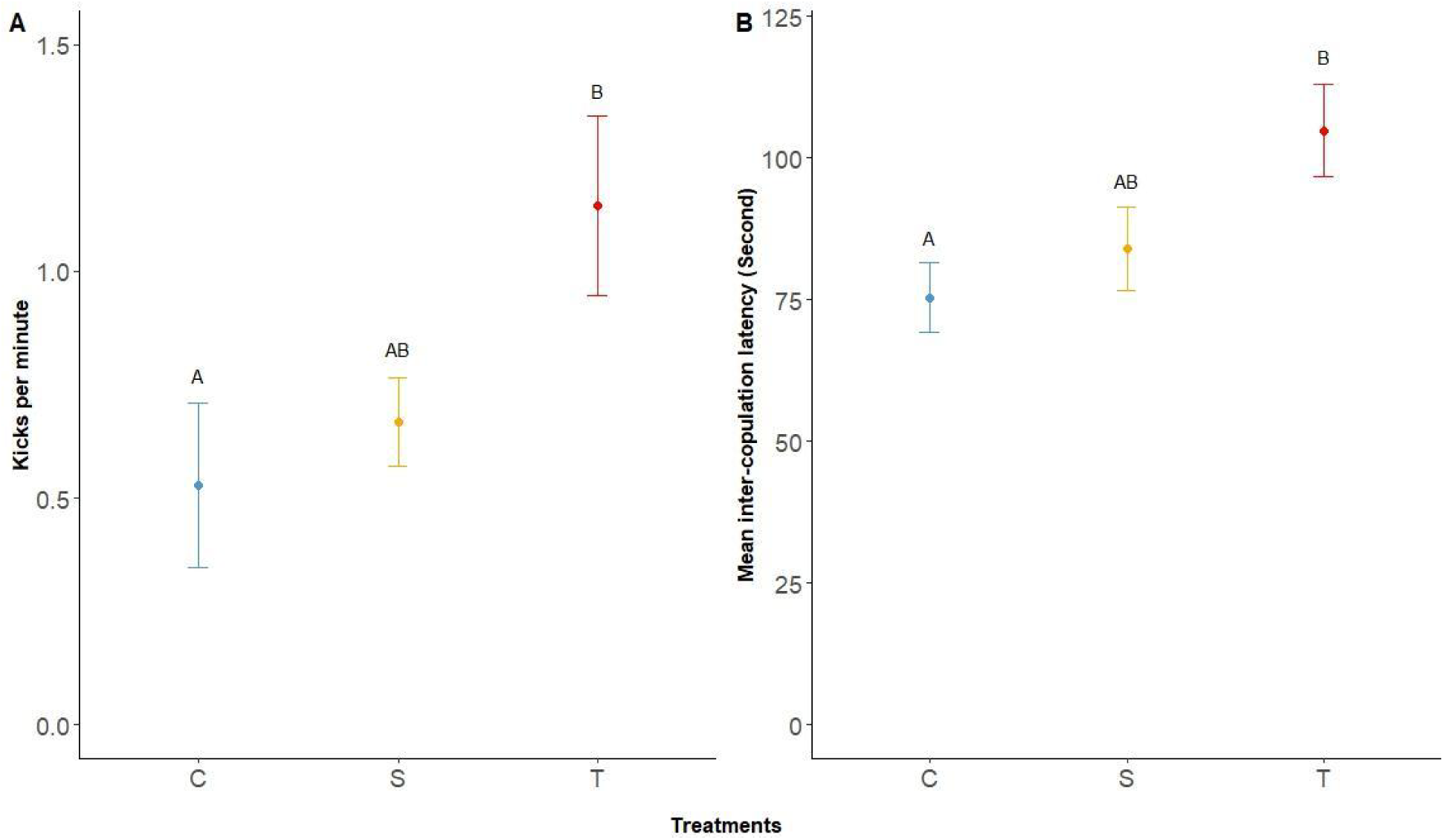
Kicks received and mean inter-copulatory latency of the Control (C), Sham (S) and hind leg tarsi ablated (T) males. Treatments not sharing the same letter are statistically different.

## Discussion

Both in the laboratory controlled condition as well as in semi-natural conditions, Parthenium beetles appeared to show an inordinately long mating. Afaq & Omkar (2017) and Bhaisare et al. (2021) reported average mating duration of 4.33 hours and 240.34 minutes respectively. In our study, the mating associations were similarly long, 322.30 and 235.35 minutes on an average in the laboratory and semi-natural setups, respectively. Interestingly, our observation suggests that this extraordinarily long mating is not a single copulation, but a series of copulations within a single mount. Each mount was found to be a set of complex behaviours including copulation, leg rubbing and latent inter-copulatory phases. Copulations usually continue almost for the entire duration of the mounting and, thus, should not be labelled “post-copulatory mate guarding” as suggested by Bhaisare et al. (2021). Moreover, the mounting duration showed high variability in our study, possibly reflecting the natural variation present in the population. To the best of our knowledge, this is the first study to report this copulatory pattern in this species. Though not unique, such mating behaviour is rather rare across animal taxa. In the following section, we provide a brief overview of the literature on such mating behaviour across animal taxa, and discuss our finds in light of sexual selection theories.

Dickinson (1986) classified prolonged mating associations in insects into four categories. First, a brief copulatory phase followed by a long post-copulatory passive/guarding phase (e.g., Saeki et al., 2005). Second, a long passive/guarding phase followed by a brief copulation (e.g., Parker & Vahed, 2010; Bennett et al., 2012). Third, a long copulatory phase in which genital contact is maintained in almost the entirety of the prolonged association (e.g., Wang et al., 2008; Yoshizawa et al., 2014). Fourth, repeated copulation with the same female and showing courtship or aggression or riding passively between intromissions. Mating behaviour observed in the Parthenium beetles seems to correspond to the fourth category. This behaviour has two characteristic elements – long mating association and repeated copulations. These two elements might not be independent of each other, as repeated copulations extend the mating duration. Evidence of such reproductive behaviour in insects has been rare (see Wright, 1960; Hagley, 1965; Alcock, 1976; Smith, 1979; Hughes, 1981; Johnson, 1982; Wickler & Seibt, 1985; Dickinson, 1986; Eberhard et al., 1993; Otronen, 1994; Hanks et al., 1996; Cueva Del Castillo et al., 1999; Cueva Del Castillo, 2003; Suzaki & Miyatake, 2011), with no specific taxonomic distribution pattern.

There are two broad theories by which such mating behaviour can be explained – (a) fertility and (b) sexual selection (O’Donald, 1978; Suter & Parkhill, 1990; deCatanzaro, 1991; Dixson, 1995; Eberhard, 1997; Hirai & Kimura, 1999; Omkar et al., 2006). These two theories are not mutually exclusive. Our results suggest that repeated copulations have both fertility as well as competitive ability consequences.

When there is a linear relationship between mating duration and sperm transfer by males (Thornhill, 1976; Laird et al., 2004), and a positive correlation between fertility and the number of copulations; repeated copulation and long mating can be adaptive. The latter is possible if the number of sperm transferred in each copulation is insufficient to fertilize the available eggs (Hunter et al., 1993; Cook, 1999). We found, in *Z. bicolorata*, the number of sperm transferred to be significantly lower if only a single copulation was allowed compared to if 15 or higher number of copulations were allowed. Clearly, the initial copulations (of the entire episode of ∼55 copulations that constitute a full mounting) are needed in this species to ensure the transfer of sufficient sperm. This inference was further upheld by our fecundity and hatchability results from the same interrupted mating assay, wherein we found these two measures of fertility of the females that went through only a single copulation to be significantly lower compared to those that were allowed 15 or higher number of copulations. The exact minimum number of copulations needed to ensure optimal fertility could not be deduced from our results due to the design of the interruption treatment. However it is amply clear that beyond this minimum sufficient number of copulations, further copulations are perhaps not needed to ensure fertility per se. Importantly, our observations are consistent with previous reports, where mating interruption has been done based on duration instead of number of copulations (Afaq & Omkar, 2017; Bhaisare et al., 2021).

Our results further indicate that such copulatory behaviour could impact competitive ability of the males through the modulation of female behaviour. Experimentally induced shorter mounting, i.e., lower number of copulations resulted in a significantly longer subsequent mounting (i.e., effectively remating) in the experimental females. Though we could not measure paternity consequence of such alteration of female behaviour, it is not unreasonable to consider this as an indication of the putative role of repeated copulations in paternity assurance. In many insects, seminal fluid proteins (products of the male accessory gland) modulate female remating behaviour (Radhakrishnan & Taylor, 2007; Yamane & Miyatake, 2012; Gabrieli et al., 2016). One of the best studied examples is fruit flies, *Drosophila melanogaster,* wherein numerous seminal fluid proteins have been identified to have roles such as, reduction of female remating tendency, increase in foraging activity, increase in egg production etc. (See Chapman & Davies, 2004; Avila et al., 2011 for reviews). Interestingly, only a few minutes following the initiation of a copulation is usually sufficient for sperm transfer, while the remaining copulatory period is needed for the transfer of seminal fluid components other than sperm (Gilchrist & Partridge, 2000). Hence, variation in copulation duration is thought to represent variation in the transfer of seminal fluid proteins – resulting in variation in sperm competitive ability of the males in this species (Wigby et al., 2009; Dhole & Servedio, 2014; Patlar & Ramm, 2020, Patlar & Civetta, 2022). In our investigation, since we do not have the means of quantifying paternity success, we are unable to extend our observation on female subsequent mounting stated above. Nonetheless, if mounting duration in our beetles is equivalent to the duration of copulation in other insects including fruit flies, our results might be the first to connect long mounting and post-copulatory competitive ability in Parthenium beetles.

Besides fertility and competitive ability, long mating could also be linked to mate quality assessment, especially if the males transfer nutrients during mating (Bloch Qazi et al., 1996; Bonduriansky, 2001; Fedina & Lewis, 2007). However, in our investigation, there was no evidence of such nutrient transfer either in the form of decreased starvation resistance or reduced egg provisioning upon mounting interruption. On the contrary, interrupted mounting improved starvation resistance in both sexes – indicating that long mounting is costly. Importantly, starvation survival time, as measured in our assay, measured both starvation as well as desiccation resistance. Given the methods, it is not possible to separate out the starvation resistance from desiccation effect. Nonetheless, such cost may come from missed foraging opportunities during mounting and/or the energetic requirement of repeated copulation. Further, for females, the mounting cost can also represent sexual antagonism (Magurran and Seghers, 1994; Stockley, 1997; Jones et al., 2010), a possibility which needs further exploration.

In summary, we found some subtle evidence to suggest that the long mounting observed in *Z. bicolorata* has an important role in facilitating not only sperm transfer, but also non-sperm components of seminal fluid. In addition, we also showed for the first time that such long mounting could be connected to modulation of females’ subsequent mating behaviour. In addition, long mounting in these beetles can also be the copulatory mate guarding, which helps males physically guard their mates. Guarding polyandrous females, when the risk of sperm competition is high, is very common in insects (Caroll, 1991; Alcock, 1994; Chaudhary & Mishra, 2015). However, guarding could have been achieved by engaging in single copulation for a prolonged period instead of copulating repeatedly. Though convenient and widely accepted, we advocate treating mate guarding as a hypothesis which needs to be robustly tested, and is non-mutually exclusive to other theories as copulatory courtship.

One of the most interesting findings of our study was the serendipitous observation of ‘kicking’ from females. Beetles with truncated hind legs received more kicks per minute from the females compared to control and sham females. Similarly, the mean inter-copulatory latency was also significantly higher in males with truncated hind legs. Both results are suggestive of leg rubbing to function in reducing the female resistance. However, ablation did not affect the mounting latency, mean copulation duration, and mounting duration, contrary to our prediction. Combining our data on re-mounting duration and leg rubbing suggests that the behaviours involved alongside copulations may have additional roles. Increased inter-copulation latency and increased kicking in hind leg tarsi ablated males clearly demonstrate the alteration in female behaviours in response to the male behaviours in the inter-copulatory phase. Performing leg rubbing and associated behaviours for a prolonged time could be energy-intensive, as indicated in our starvation resistance data. Thus, the ability to perform leg rubbing and associated behaviours for a prolonged time could be an honest signal of male genetic quality (Tallamy et al., 2003). Females may differentially transport, store, and utilize the sperm of different males based on the behaviours performed by the males, which ultimately determine male fertilization success (Bloch Qazi, 2003; Edvardsson & Arnqvist, 2005). We suggest that leg rubbing in the Parthenium beetle is a copulatory courtship behaviour that may have evolved in response to sexual selection by cryptic female choice.

Whether copulatory courtship in the form of leg rubbing truly reflects the male quality would add insights into our present understanding of long mounting involving repeated copulations. It would also be interesting to check the change in sperm precedence and paternity share in response to the changes in mounting duration/number of copulations and stimulation of the females by leg rubbing. Since the longer mating association did not improve female fitness, it is important to know whether there is sexual conflict over optimal duration of mating association and number of copulations. If sexual conflict holds group-level consequences (Eldakar & Gallup, 2011), it may affect the local population growth, which would in turn reduce the biocontrol efficiency of these beetles.

## Acknowledgement

We thank Ranjit Kumar Sahoo for his guidance in establishing the beetle population and initial standardization of population maintenance protocols. We are grateful to Purbasha Dasgupta for her comments and suggestions on an earlier version of the manuscript. Further. the authors acknowledge Tanya Verma, Purbasha Dasgupta, Subhasish Halder, Usha Kiran Sahoo, Gayathri Pramod, Anish Koner, Gokul Bhaskaran, Debapriya Dari, Anitta Cherian and Shramana Kar for their help in the experiments. RSP thanks Indian Institute of Science Education and Research Berhampur for financial support in the form of Junior and Senior Research Fellowship.

