## Supplementary information for "Long mounting with repeated copulations in *Zygogramma bicolorata*: A test of adaptive significance of a curious mating behaviour"

Mating behaviour under semi-natural conditions:

Beetles from the F2 generation were collected as virgins in single-sex adult boxes. Beetles were acclimated to outdoor conditions for 1, 2, 4, 8, 12, 12 and 12 hours respectively (starting at 6 am on each day) on seven consecutive days prior to the day of observation. On the 12^th^ day, randomly picked virgin females and males were released in close proximity on the Parthenium plants grown in pots. Generally, the males mounted almost immediately once they encountered the females. If they did not interact or moved apart from each other, the males were released back into the proximity of the females. New pairs were tried if the pairs did not remain in proximity. Each observer observed only one pair at any given time. A total of 20 observations were staggered over seven days to accommodate the logistic issues. The observation started around 10 a.m. each day and continued until the end of the mounting. Similar to laboratory observation, release time, mounting time, beginning and end time of each copulation, and additional behaviours were noted.

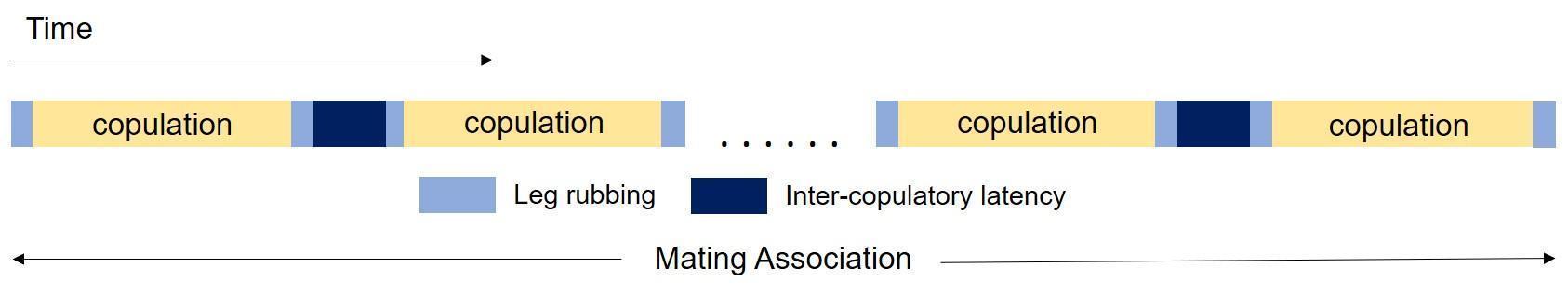

**Figure S1:** Schematic representation of the behaviours displayed during a mating association. The mating association continues for an extended period which consists of a series of copulations. Each copulation is preceded and followed by leg rubbing. Two copulations are punctuated by an inter-copulatory latent period.

**Table S1:** Descriptive statistics of the behavioural parameters measured during the characterization of the mating behaviour. Values indicate mean durations and standard errors (SE) in minutes from observations in laboratory and semi-natural conditions.

| **Parameter** | **Laboratory Condition** | | **Semi-natural Condition** | |
| --- | --- | --- | --- | --- |
|  | **Mean (minute)** | **SE** | **Mean (minute)** | **SE** |
| Mounting latency | 38.64 | 5.5 | 1 | 0.22 |
| Copulation latency | 0.98 | 0.26 | 0.35 | 0.12 |
| First copulation duration | 12.22 | 0.83 | 6.11 | 0.42 |
| Last copulation duration | 8.89 | 0.68 | 6.28 | 0.77 |
| Mean copulation duration | 3.95 | 0.16 | 2.67 | 0.11 |
| Mean inter-copulation latency | 1.52 | 0.11 | 1.31 | 0.15 |
| Mounting duration | 322.3 | 24.31 | 235.35 | 35.46 |
| Number of copulations | 55.5 | 5.04 | 56.75 | 9.24 |

**Table S2:** Results of the Wilcoxon signed-rank test showing comparisons between first, last, and mean copulation duration in laboratory and semi-natural setup. Statistically significant *p*-values are highlighted in boldface.

| **Contrast** | **Laboratory Condition** | | **Semi-natural Condition** | |
| --- | --- | --- | --- | --- |
|  | **V** | ***p*-value** | **V** | ***p*-value** |
| First-Mean | 1023 | **<0.001** | 153 | **<0.001** |
| Last-Mean | 974 | **<0.001** | 158 | **<0.001** |

**Table S3:** Results of pairwise comparison between interruption treatments in sperm transfer. Statistically significant *p*-values are highlighted in boldface.

| **Contrast** | **Estimate** | **SE** | **df** | **t ratio** | ***p*-value** |
| --- | --- | --- | --- | --- | --- |
| C1-C15 | 2.74$\times$10^-4^ | 6.65$\times$10^-5^ | 44 | 4.117 | **0.001** |
| C1-C30 | 2.95$\times$10^-4^ | 6.60$\times$10^-5^ | 44 | 4.472 | **<0.001** |
| C1-C50 | 2.97$\times$10^-4^ | 6.60$\times$10^-5^ | 44 | 4.507 | **<0.001** |
| C1-Control | 2.93$\times$10^-4^ | 6.61$\times$10^-5^ | 44 | 4.426 | **<0.001** |
| C15-C30 | 2.13$\times$10^-5^ | 1.14$\times$10^-5^ | 44 | 1.873 | 0.346 |
| C15-C50 | 2.35$\times$10^-5^ | 1.12$\times$10^-5^ | 44 | 2.1 | 0.238 |
| C15-Control | 1.87$\times$10^-5^ | 1.18$\times$10^-5^ | 44 | 1.576 | 0.520 |
| C30-C50 | 2.15$\times$10^-6^ | 7.59$\times$10^-6^ | 44 | 0.283 | 0.998 |
| C30-Control | -2.67$\times$10^-6^ | 8.54$\times$10^-6^ | 44 | -0.313 | 0.998 |
| C50-Control | -4.82$\times$10^-6^ | 8.26$\times$10^-6^ | 44 | -0.584 | 0.977 |

**Table S4:** Results of pairwise comparison between treatments in cumulative fecundity. Statistically significant *p*-values are highlighted in boldface.

| **Contrast** | **Estimate** | **SE** | **df** | **t ratio** | ***p*-value** |
| --- | --- | --- | --- | --- | --- |
| C1-C15 | -150.9 | 41.4 | 54 | -3.642 | **0.005** |
| C1-C30 | -135.1 | 41.4 | 54 | -3.262 | **0.016** |
| C1-C50 | -120.5 | 41.4 | 54 | -2.91 | **0.040** |
| C1-Control | -130.6 | 41.4 | 54 | -3.153 | **0.021** |
| C15-C30 | 15.8 | 40.5 | 54 | 0.389 | 0.995 |
| C15-C50 | 30.3 | 40.5 | 54 | 0.749 | 0.944 |
| C15-Control | 20.2 | 40.5 | 54 | 0.5 | 0.987 |
| C30-C50 | 14.6 | 40.5 | 54 | 0.36 | 0.996 |
| C30-Control | 4.5 | 40.5 | 54 | 0.111 | 1.000 |
| C50-Control | -10.1 | 40.5 | 54 | -0.249 | 0.999 |

**Table S5:** Results of pairwise comparison between treatments in mean hatchability. Statistically significant *p*-values are highlighted in boldface.

| **Contrast** | **Estimate** | **SE** | **df** | **z ratio** | ***p*-value** |
| --- | --- | --- | --- | --- | --- |
| C1-C15 | -0.616 | 0.148 | Inf | -4.153 | **<0.001** |
| C1-C30 | -0.611 | 0.145 | Inf | -4.204 | **<0.001** |
| C1-C50 | -0.658 | 0.148 | Inf | -4.442 | **<0.001** |
| C1-Control | -0.519 | 0.146 | Inf | -3.555 | **0.003** |
| C15-C30 | -0.005 | 0.137 | Inf | 0.040 | 1.000 |
| C15-C50 | -0.042 | 0.140 | Inf | -0.299 | 0.998 |
| C15-Control | -0.098 | 0.138 | Inf | 0.706 | 0.955 |
| C30-C50 | -0.047 | 0.137 | Inf | -0.345 | 0.997 |
| C30-Control | 0.092 | 0.135 | Inf | 0.682 | 0.960 |
| C50-Control | 0.139 | 0.138 | Inf | 1.010 | 0.851 |

**Table S6:** Results of pairwise comparison between treatments in the duration of re-mounting. Statistically significant *p*-values are highlighted in boldface.

| **Contrast** | **Estimate** | **SE** | **df** | **t ratio** | ***p*-value** |
| --- | --- | --- | --- | --- | --- |
| C30-C50 | 39.8 | 34.4 | 51 | 1.156 | 0.485 |
| C30-Control | 151.1 | 34.9 | 51 | 4.324 | **<0.001** |
| C50-Control | 111.3 | 35.4 | 51 | 3.144 | **0.008** |

Effect of time of the day on mating behaviour:

In Experiment 2, remating observations started at different times of the day, which may possibly influence the mating behaviour. To test for any effect of time of the day on mating, we conducted a parallel experiment to the remating assay. We set up mating of 10 pairs approximately at the same time the rematings for each treatment were set. Any difference in the mounting latency, and mounting duration were checked.

Negative binomial Generalized Linear Model (GLM) was used to analyze mounting latency data using the MASS package (Venables & Ripley, 2002). Mounting duration data was fit into a linear model. Copulation latency was not analyzed since 19 out of 26 observations had zero values. Time of the day was incorporated as the fixed factor in the models.

We did not find any significant effect of time of the day on mounting latency (*F*_2,26_=0.024, *p*=0.976) and mounting duration (*F*_2,27_=0.056, *p*=0.946). Therefore, the observed effect of interruption in the first mounting affecting the re-mounting duration is unlikely to be confounded by any effect of time of the day.

**Table S7:** Results of the cox proportional hazard analysis on survivorship of females and males (combined) under starvation that copulated till C1, C30 and uninterrupted (Control). Hazard rates are expressed relative to the hazard rate of default level of each fixed factor constrained to be 1. The default level of treatment is ‘Virgin’, while that of sex is ‘Female’. Lower CI and Upper CI indicate lower and upper bounds of 95% confidence intervals. Confidence intervals that do not contain 1 signify statistical significance.

| **Fixed coefficients** | **Hazard ratios** | **Lower CI** | **Upper CI** | **z** | ***p-value*** |
| --- | --- | --- | --- | --- | --- |
| Treatment: C1 | 1.403 | 0.801 | 2.455 | 1.185 | 0.236 |
| Treatment: C30 | 1.985 | 1.130 | 3.488 | 2.384 | **0.017** |
| Treatment: Control | 3.224 | 1.743 | 5.961 | 3.732 | **<0.001** |
| Sex: Male | 0.869 | 0.495 | 1.527 | -0.487 | 0.626 |
| Treatment C1 × Sex Male | 1.540 | 0.694 | 3.417 | 1.061 | 0.288 |
| Treatment C30 × Sex Male | 1.363 | 0.616 | 3.017 | 0.764 | 0.445 |
| Treatment Control × Sex Male | 1.185 | 0.525 | 2.671 | 0.408 | 0.683 |

**Table S8:** Pairwise comparison table showing differences between treatments in their survival under starvation. Statistically significant *p*-values are highlighted in boldface.

|  | **Virgin** | **C1** | **C30** |
| --- | --- | --- | --- |
| **C1** | **0.043** |  |  |
| **C30** | **0.003** | 0.098 |  |
| **Control** | **<0.001** | **<0.001** | **0.036** |

Effect of interruption treatment on the egg size:

Results of the analysis revealed a significant effect of interruption treatment on egg size (Table 1). C15 and C50 females laid larger eggs compared to C1 and Control (Figure S2, Table S9). Eggs from the C30 treatment group were the smallest compared to all other treatments (Figure S2, Table S9).

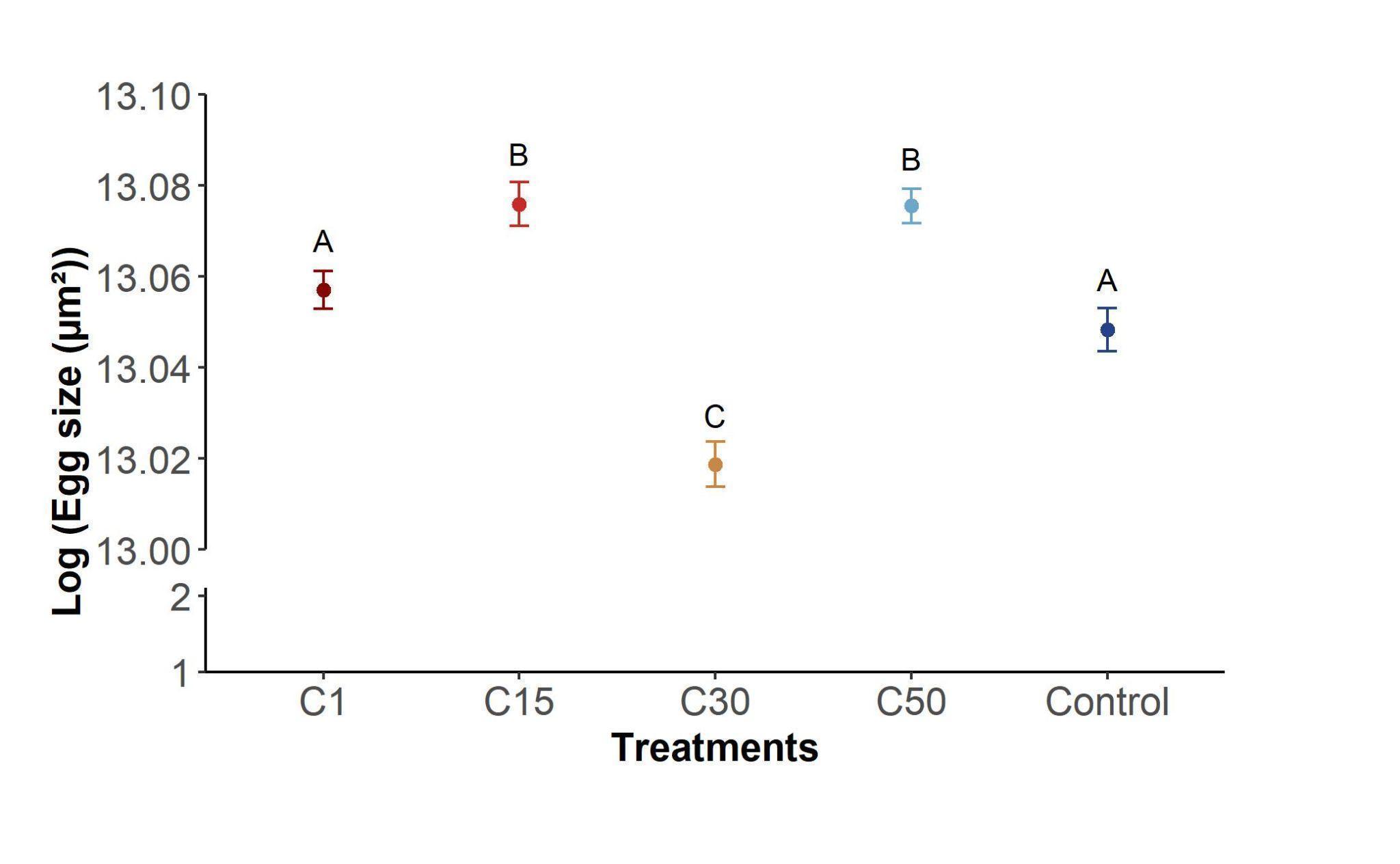

**Figure S2:** Sizes of eggs laid by females mounted till copulation 1 (C1), 15 (C15), 30 (C30), 50 (C50), or uninterrupted (Control) displayed in the logarithmic scale. Egg sizes in the plot represent pooled data from day 1, 5, 10 followed by 10 day intervals till day 60 post-mating. The circle and the error bar represent the mean and standard error, respectively. Treatments not sharing the same letter are statistically different.

**Table S9:** Results of pairwise comparison between treatments in the egg size. Statistically significant *p*-values are highlighted in boldface.

| **Contrast** | **Estimate** | **SE** | **df** | **t ratio** | ***p*-value** |
| --- | --- | --- | --- | --- | --- |
| C1-C15 | 4.16$\times$10^-8^ | 1.35$\times$10^-8^ | 1347 | 3.087 | **0.018** |
| C1-C30 | -8.12$\times$10^-8^ | 1.39$\times$10^-8^ | 1347 | -5.831 | **<0.001** |
| C1-C50 | 3.84$\times$10^-8^ | 1.33$\times$10^-8^ | 1347 | 2.878 | **0.033** |
| C1-Control | -1.76$\times$10^-8^ | 1.38$\times$10^-8^ | 1347 | -1.275 | 0.707 |
| C15-C30 | -1.23$\times$10^-7^ | 1.37$\times$10^-8^ | 1347 | -8.976 | **<0.001** |
| C15-C50 | -3.25$\times$10^-9^ | 1.31$\times$10^-8^ | 1347 | -0.249 | 0.999 |
| C15-Control | -5.92$\times$10^-8^ | 1.35$\times$10^-8^ | 1347 | -4.372 | **<0.001** |
| C30-C50 | 1.20$\times$10^-7^ | 1.35$\times$10^-8^ | 1347 | 8.833 | **<0.001** |
| C30-Control | 6.36$\times$10^-8^ | 1.40$\times$10^-8^ | 1347 | 4.551 | **<0.001** |
| C50-Control | -5.59$\times$10^-8^ | 1.34$\times$10^-8^ | 1347 | -4.177 | **<0.001** |

The number of copulations was found to influence the egg size. Females that copulated only once laid similar sized eggs compared to the control females, but smaller compared to the females that copulated 15 and 50 times. Higher egg sizes in C15 and C50 females could be due to higher egg provisioning following utilization of nuptial gifts from the male. Surprisingly, egg sizes in control females were similar to the C1 females and significantly lower than the C15 and C50 females. The nutrient donations may not be utilized in reproduction if the cost of mating for an extended time is significant. Resource utilization in egg provisioning could hence be reduced, resulting in smaller egg sizes in control females. Although our starvation resistance data reveal some cost of mating in C50 females in terms of survival under starvation, this cost paid by C15 and C50 females could be insufficient to reduce the egg provisioning. The egg sizes can also change due to egg number and size trade-offs (Gibbs et al., 2005). Our data on fecundity from earlier generation, however, does not support this possibility. Why egg sizes in C30 females were smaller than all other treatments also remains a puzzle to us.

**Table S10:** Results of pairwise comparison between ablation treatments on the number of kicks received per sampled minute by the males. Statistically significant *p*-values are highlighted in boldface.

| **Contrast** | **Estimate** | **SE** | **df** | **t ratio** | ***p*-value** |
| --- | --- | --- | --- | --- | --- |
| C - S | -0.138 | 0.24 | 31 | -0.575 | 0.835 |
| C - T | -0.616 | 0.235 | 31 | -2.619 | **0.035** |
| S - T | -0.478 | 0.235 | 31 | -2.032 | 0.121 |

**Table S11:** Results of pairwise comparison between ablation treatments on the mean copulation latency. Statistically significant *p*-values are highlighted in boldface.

| **Contrast** | **Estimate** | **SE** | **df** | **t ratio** | ***p*-value** |
| --- | --- | --- | --- | --- | --- |
| C-S | -8.71 | 11.2 | 27 | -0.775 | 0.721 |
| C-T | -29.50 | 10.5 | 27 | -2.807 | **0.024** |
| S-T | -20.79 | 10.5 | 27 | -1.978 | 0.137 |
